# Stress-responsive transcription factor families are key components of the core abiotic stress response in maize

**DOI:** 10.1101/2025.02.15.638452

**Authors:** Anna C.H. Pardo, Jeremy D. Pardo, Robert VanBuren

## Abstract

Abiotic stresses, including drought, salt, heat, cold, flooding, and low nitrogen, have devastating impacts on agriculture and are increasing in frequency globally due to climate change. Plants can experience multiple abiotic stresses simultaneously or sequentially within a single growing season, and combinatorial stresses elicit shared or overlapping molecular and physiological responses. Here, we searched for core stress responsive genes in maize across diverse abiotic stressors through meta-analysis of public RNAseq data. Our analysis revealed significant heterogeneity in gene expression across datasets due to factors such as tissue type, genotype, and experimental conditions, which we mitigated through batch correction. Using nearly 1,900 RNAseq samples with both traditional set operations and a novel random forest machine learning approach, we identified a core set of 744 stress-responsive genes across the six stresses. These core genes are enriched in transcription factors, including stress-responsive families such as AP2/ERF-ERF, NAC, bZIP, HSF, and C2C2-CO-like. Co-expression network analysis demonstrated that these core TFs are co-expressed with stress-specific peripheral genes, supporting their role in regulating both generalized and stress-specific responses. Our results suggest that maize employs a conserved yet flexible transcriptional strategy to respond to abiotic stresses, with core TFs acting as potential regulators of both universal and stress-specific pathways. These findings provide a valuable resource for understanding stress tolerance mechanisms and for guiding future breeding and engineering efforts to enhance maize resilience under climate change.

## INTRODUCTION

Abiotic stresses such as drought, salt, flooding, heat, cold, and low nitrogen can have severe impacts on agricultural crops. Climate change has slowed the growth of agricultural productivity in the last half century, and events such as droughts and floods are projected to increase in frequency under our dynamic climate (Calvin et al., 2023). This is expected to cause a general decrease in global crop productivity (Raza et al., 2019). Maize (*Zea mays*) is one of the most important food, feed, and biofuel crops globally, and like all crops, may experience many abiotic stressors of different types individually or in combination throughout its growing season. Thus, understanding the basis of abiotic stress response on the molecular level is important for eventual improvement of maize stress tolerance and increased agricultural resilience in the face of climate change.

Abiotic stresses present many challenges to plants at the cellular and molecular levels. For example, most abiotic stressors, including cold, low nitrogen, drought, and flooding, are known to inhibit photosynthesis (Singh and Thakur, 2018; Li et al., 2019; Wu et al., 2019; Qi et al., 2021). Typically this leads to a buildup of reactive oxygen species, which can damage cell components such as membranes. Antioxidants such as glutathione, which scavenge excess reactive oxygen species, are instrumental in the cellular response to stresses including salt, drought, flooding, and cold (Aslam et al., 2021). In stresses with an osmotic component, such as drought, salt, and cold, it is common for plants to accumulate osmolytes/compatible solutes such as sugars, amino acids including proline and glycine betaine, and polyamines; these molecules function as protectants against water loss and may also have antioxidant activity (Sharma et al., 2019). Polyamines are also hypothesized to act as plant growth regulators (EL Sabagh et al., 2022), and both transformation for increased polyamine production and exogenous polyamine application are known to improve plant stress tolerance (Minocha et al., 2014). Many of these responses are mediated by phytohormones such as ethylene, abscisic acid (ABA), and jasmonic acid (JA), and are similar across different stressors. For example, ethylene and JA are noted as active regulators under various stress conditions, including high and low temperature, drought, and salt among others (EL Sabagh et al., 2022).

Transcription factors (TFs) regulate stress responses in both phytohormone-dependent and independent manners. For example, TFs in the ABF subfamily of the bZIP family are activated by signal transduction following ABA recognition; following their activation, they go on to activate the expression of NAC and AP2/ERF family TFs (Yoon et al., 2020). These TFs then go on to regulate other stress-responsive genes (Mizoi et al., 2012; Nakashima et al., 2012; Shao et al., 2015), which act to impact the plant’s response to a given stress. Given the similarity of molecular and cellular responses to different stresses, it is reasonable to hypothesize the existence of a core stress response controlled by core stress-responsive genes.

Generally, core stress responses have been studied using two approaches: the first involves gathering transcriptomic data from an experiment with multiple different stress treatments; the second, conducting a meta-analysis of previously published transcriptomic data, either from microarrays or RNA-sequencing. Meta-analyses or multi-stressor experiments have previously been conducted in cotton (Tahmasebi et al., 2019), rice (Cohen and Leach, 2019), sesame (Dossa et al., 2019), *Brassica napus* (Zhang et al., 2019), and *Arabidopsis thaliana* (Sanchez-Munoz et al., 2024; Shintani et al., 2024), as well as maize (Li et al., 2017), among others. These previous meta-analyses often use a relatively limited number of studies and sometimes only one per stressor, which may limit the power of the meta-analysis to draw biologically relevant conclusions. All meta-analyses cited here re-analyzed data from at most five hundred transcriptome samples. The largest of these, by (Sanchez-Munoz et al., 2024), re-analyzed 500 samples from 23 different studies, including both microarray and RNA-seq data, from a total of 11 stressors in both roots and shoots. Most other meta-analyses examined only four or five stressors.

In the current study, we re-analyzed nearly 1,900 RNA-sequencing samples from 39 different maize stress experiments, spanning a wide variety of genotypes, growth environments, tissues, and developmental stages. We leveraged these data to identify core genes using both a standard set operations approach, which classified genes as typically up- or downregulated across stress conditions, and a random forest classification approach, which selected genes with the highest feature importance. As part of our set operations, we also identified stress-specific, or stress-specific, genes. Analysis of the core genes revealed that they were enriched in several stress-related TF families, which were found to be coexpressed both with other core genes and with stress-specific genes, indicating their possible role in regulating not only the core response, but stress-specific responses as well.

## METHODS

### Curating maize RNAseq data

This study utilized publicly available and previously published abiotic stress RNA-sequencing (RNA-seq) data in maize from the NCBI Sequence Read Archive (SRA). Only RNA-seq data that could be linked with published papers were used. Data were collected for drought, cold, heat, salt, flooding, and low nitrogen, which are the best studied abiotic stresses in maize. Polyethylene glycol and similar treatments such as sugar alcohols were not included in the drought data. BioProjects were only included in the study if they had at least one well-documented stress time point as well as either a control treatment or samples taken at experiment initiation. Thirty-nine BioProjects containing 1,981 samples total met these criteria, and were selected for use in this study. Each of the six abiotic stresses had at least three independent experiments (BioProjects). The dataset includes stress-tolerant and sensitive maize genotypes, and hybrids, but not mutants or transgenic plants. Most samples were from leaf or other photosynthetic tissues, but roots, reproductive, and seed tissues were also included. Studies with any number of replicates were included.

### Processing the RNAseq data

All data were downloaded from the SRA using the prefetch and fasterq-dump commands from sratoolkit v2.11.2 (https://github.com/ncbi/sra-tools). The raw RNA-seq reads were processed using Nextflow v23.04.1 and the nf-core rnaseq pipeline v3.11.1 (Di Tommaso et al., 2017; Ewels et al., 2020; dependencies: https://github.com/nf-core/rnaseq/blob/master/CITATIONS.md). Within the pipeline, reads were trimmed using fastp (Chen et al., 2018) and transcripts were quantified using Salmon (Patro et al., 2017). Length-scaled TPM were then generated using tximport (Soneson et al., 2016). The exception to our processing workflow was a low nitrogen dataset (Ying et al., 2023; BioProject: PRJNA904734), which was previously analyzed and processed with STAR (Dobin et al., 2013). The convertCounts() function in the DGEobj.utils R package v1.0.6 (https://CRAN.R-project.org/package=DGEobj.utils) was used in R 4.2.1 (R Core Team, 2022) to convert the provided count matrix into TPM.

All transcripts were pseudoaligned to the B73 v5 genome (Hufford et al., 2021). Our dataset contains a diversity of maize genotypes, including multiple with chromosome scale assemblies, and we initially processed the data by pseudoaligning each genotype to the corresponding reference genome when available. However, we ultimately decided to map all data to B73 for the following reasons: 1) mapping rates were similar when pseudoaligning to B73 vs. the corresponding genome for several inbred lines (see Supplemental Table 1); 2) many genotypes do not have sequenced genomes or are hybrids; 3) the B73 reference annotation is manually curated and more complete than others; and 4) graph based annotations and syntenic gene groupings are still incomplete, making it challenging to create an informative set of comparable genes across genotypes for downstream comparisons. Thus, given the goals of this study, and the above limitations, we chose to use a single genome for read mapping.

### Data exploration of experimental factors

Data exploration was conducted with principal component analysis (PCA) using the full dataset that includes 12 tissue types (referred to as “all tissues” hereafter), and for photosynthetic tissues only, which included leaf, leaf meristem, and shoot samples. The raw gene expression (transcripts per million; TPM) values were filtered to remove genes with zero variance across samples, and the TPM values were log2 +1 transformed using numpy v1.24.3 (Harris et al., 2020). RNAseq datasets were collected using different sequencing machines, read lengths, coverage, and under different experimental conditions, and we reduced the batch effect of BioProject using pyComBat v0.3.3 (Behdenna et al., 2023). PCA was run on the uncorrected and batch corrected data for the full dataset and photosynthetic data only, and the first two principal components were calculated using scikit-learn v1.2.2 (Pedregosa et al., 2011) and plotted using matplotlib v3.7.1 (Hunter, 2007).

To further explore heterogeneity within the dataset, we modeled the first principal component (PC1) of log-transformed, non-batch corrected TPM as a function of genotype, BioProject, treatment, and tissue, using the lm() function in R v4.2.1 (R Core Team, 2022). Interaction effects were excluded for computational efficiency, and because interpreting the biological significance of interaction effects can be challenging. This same modeling approach was repeated for PC1 of batch corrected TPM, allowing us to compare results between the two models.

### Differential expression analysis and identification of stress-induced genes

Changes in gene expression between stressed and control samples were evaluated using two different methodologies: differential expression analysis and fold change or the ratio of gene expression in stress-treated (T) and control, or non-treated (N) samples (TN-ratio; described in detail below) (Shintani et al., 2024). For differential expression analysis, tximport (Soneson et al., 2016) was used to generate the raw gene expression values (in TPM), and DESeq2 v1.38.3 (Love et al., 2014) was used to identify statistically differentially expressed genes. Differential expression was calculated as a function of an identifier containing genotype, treatment, time point, developmental stage, and tissue information for each sample. The default model was used. Only sets of samples containing at least 3 replicates were used for differential expression. TN-ratio was calculated using the formula from (Shintani et al., 2024) as follows:

TN-ratio = (stress-treated TPM+1)/(non-treated TPM+1)

For our study, TN-ratio was calculated on a per-experiment basis where for a given BioProject, the mean TPM was calculated for each sample and/or replicate within the control or stress treated groups, and this was used to calculate TN-ratio.

We used criteria as outlined in (Shintani et al., 2024), where genes with a TN-ratio of greater than 2 were considered upregulated under the respective abiotic stress, and those with a TN-ratio of less than 0.5 were considered downregulated under stress. For set operations, we calculated the union of upregulated and downregulated genes for each experiment. The core genes shared across all six stressors were identified as the intersection of these sets, representing genes consistently differentially expressed under all stresses. Genes that were only differentially expressed for a subset or only one stress condition were also identified, and we refer to these sets as ‘stress-specific genes’. The overlap of up- and downregulated genes among experiments within each stressor was also examined.

This analysis and others subsequently described in this section were initially conducted both on all samples in the dataset and on samples from photosynthetic tissues only. However, while there were some differences in numbers of core and stress-specific genes between all tissues and photosynthetic tissues, there were few differences between tissue sets in the quality of random forest modeling (described below), and downstream analyses yielded results of interest only for all tissues. Thus, this paper focuses mainly on results found with all tissues.

### Hierarchical clustering of abiotic stresses

To determine the relationships of transcriptomic responses to different abiotic stresses, we performed hierarchical clustering. BioProject-corrected, log2+1 transformed TPM values were scaled to a z-score using scikit-learn. For each of the seven treatments (six stress conditions and control), a mean expression value of the scaled and transformed TPM data was calculated and used as input for hierarchical clustering and dendrogram visualization using scipy v1.10.1 (Virtanen et al., 2020).

### Random forest binary classification

Random forest models were used to classify whether samples were stressed or control. To avoid data leakage, all stressed and associated control samples for a single stressor were held out for use as the test set. Given the hypothesized existence of a core stress response transcriptome, a random forest model tested on a stress it was not trained on was hypothesized to be able to accurately classify stressed and control samples. This was repeated for all stressors, so that each stressor was used as the test set once, resulting in a total of six models, each with separately tuned hyperparameters. Hyperparameters tuned for each model included bootstrapping, maximum tree depth, maximum features, minimum number of samples per leaf, minimum samples split, and number of estimators. In each iteration, all other samples were used for the training set.

The BioProject-corrected and log2-transformed TPM were used as features in the model such that each feature was a maize gene. SMOTE was used for upsampling to balance numbers of control vs. stress using training data. As stated above, hyperparameters were tuned separately for each model, with individual models having different stressors as the test set. The optimal hyperparameters were then used for training and making predictions.

For each of the six models fit for each set of samples, feature importance was calculated for all features (genes) used in the model. For each model, evaluation of possible core gene sets was conducted via iterative feature selection, as follows. The corrected TPM were filtered to only the top X features, where X=50, 100, 250, 500, 1,000, 1,500, 2,000, 2,500, 3,000, 4,000, 5,000, 6,000, 7,000, 8,000, 10,000, or 15,000). Following TPM subsetting, a new RF model was run on each subset and the model performance metrics accuracy, AUC, and F1 were calculated for each model. Based on the optimum model performance using the smallest subset of features, the top 6,000 most important features were extracted from each of the six individual stressor models. The intersection of these six sets of 6,000 genes each was calculated to get the core stress genes from random forest.

### Coexpression network analysis

A coexpression network was constructed with the corrected TPM of the full maize dataset using Weighted Gene Coexpression Network Analysis (WGCNA) (Langfelder and Horvath, 2008; Langfelder and Horvath, 2012). A soft threshold of 9 was used for network construction. Hub genes, i.e. genes that had a high positive or negative correlation with most genes in the same module, were identified using the module membership generated with the signedKME() function from WGCNA. A threshold value of approximately 0.86, or the 95th percentile of the absolute values of module membership, was used as the cutoff for hub gene identification. Only genes with module membership >0.86 were considered hub genes. We then found which hub genes were found in each core gene set, and used Fisher’s exact test implemented in Python using scipy.stats as described above, to test whether there were more core genes in the set of hub genes than expected by chance.

### GO term enrichment

Gene Ontology (GO) term enrichment was performed with topGO v2.50.0 (Alexa and Rahnenfuhrer, 2022) separately for Biological Process GO terms in upregulated and downregulated core genes from the set operations, random forest, and combined approaches, for both sets of tissues. We also ran GO enrichment for the stress-specific stress genes for each stressor, separately for upregulated and downregulated stress-specific sets, for each set of tissues, and for the genes in each co-expression module. Fisher’s exact test was used as the enrichment test, and the classic algorithm was used. False discovery rate (FDR) p-value correction was used to adjust p-values, and an FDR adjusted p-value of less than 0.05 was considered statistically significant. GO terms were current as of March 25, 2024.

### Transcription factor enrichment

The list of transcription factors in the *Z. mays* B73 v5 genome was downloaded from Grassius (https://grassius.org/species/Maize, Yilmaz et al., 2009). We tested for enrichment of transcription factors from all families (not any particular family) in the upregulated and downregulated core genes using a one-sided Fisher’s exact test implemented using scipy.stats v1.10.1(Virtanen et al., 2020). We also tested for transcription factor enrichment in the upregulated and downregulated stress-specific genes from each stressor. The “fdrcorrection” function from statsmodels.stats.multitest v0.13.5 (Seabold and Perktold, 2010) was used to adjust the resulting p-values. As during GO enrichment, FDR-adjusted P values less than 0.05 were considered significant.

Again using information from Grassius, the sets of all upregulated core genes and all downregulated core genes for each tissue set, from both methods combined, were tested for enrichment of each transcription factor family. TF families with FDR-adjusted P values of less than 0.05 were considered significantly enriched, while those with adjusted P values of between 0.1 and 0.05 were considered close to enriched and thus also selected for further analysis.

For the subset of core genes belonging to the enriched or near-enriched transcription factor families, we identified what co-expression network module each gene belonged to. We then used Fisher’s exact test via scipy.stats as above to determine whether modules containing these transcription factors of interest were enriched in core genes. Again, FDR was used to adjust P values and adjusted P of less than 0.05 was considered significant.

### Gene regulatory network construction and analysis

A gene regulatory network was constructed using the random forest method in GENIE3 (Huynh-Thu et al., 2010). The pyComBat-corrected, log transformed TPM data were used as input along with a list of known maize transcription factors from Grassius. Following network construction, we analyzed the differences in mean weight for target genes of core TFs from enriched families (identified above) in different gene sets, i.e. comparing core and stress-specific target genes to other target genes. This was done using Dunnett’s t test implemented in the R package DescTools. To build a distribution of p values, Dunnett’s t was repeated 5,000 times and p values for each comparison were saved. Results were considered statistically significant if the 97.5% confidence interval of the p value distribution was less than 0.05.

## RESULTS

### Exploration of maize abiotic stress gene expression data

To search for conserved molecular signatures of abiotic stress responses, we gathered published maize RNA-seq data from the NCBI Sequence Read Archive. A total of 39 BioProjects were selected, including 15 studies on drought, 8 on heat, 8 on cold, 5 on low nitrogen, 5 on salt, and 3 on flooding (Figure 1A). There were 1,872 total samples in the dataset with drought having the most samples and salt stress the fewest (Figure 1B). Most samples were collected from leaf or root tissues, though a variety of other vegetative and reproductive tissues were also represented (Table 1). Experiments were conducted in greenhouse, field, and growth chamber environments, covering developmental stages from germination to reproduction. The dataset spanned 328 maize genotypes, including both stress-sensitive and stress-tolerant lines, with inbred lines B73, W22, and Mo17 being the most frequently studied (Figure 1D). Treatment conditions varied across studies, particularly for temperature-based stress experiments. The temperature ranges defined as cold, heat, and control differed between studies, with the upper limits of cold treatments overlapping with the lower end of control conditions and a similar overlap occurring between heat treatments and control conditions (Figure 1C).

**Figure 1:**
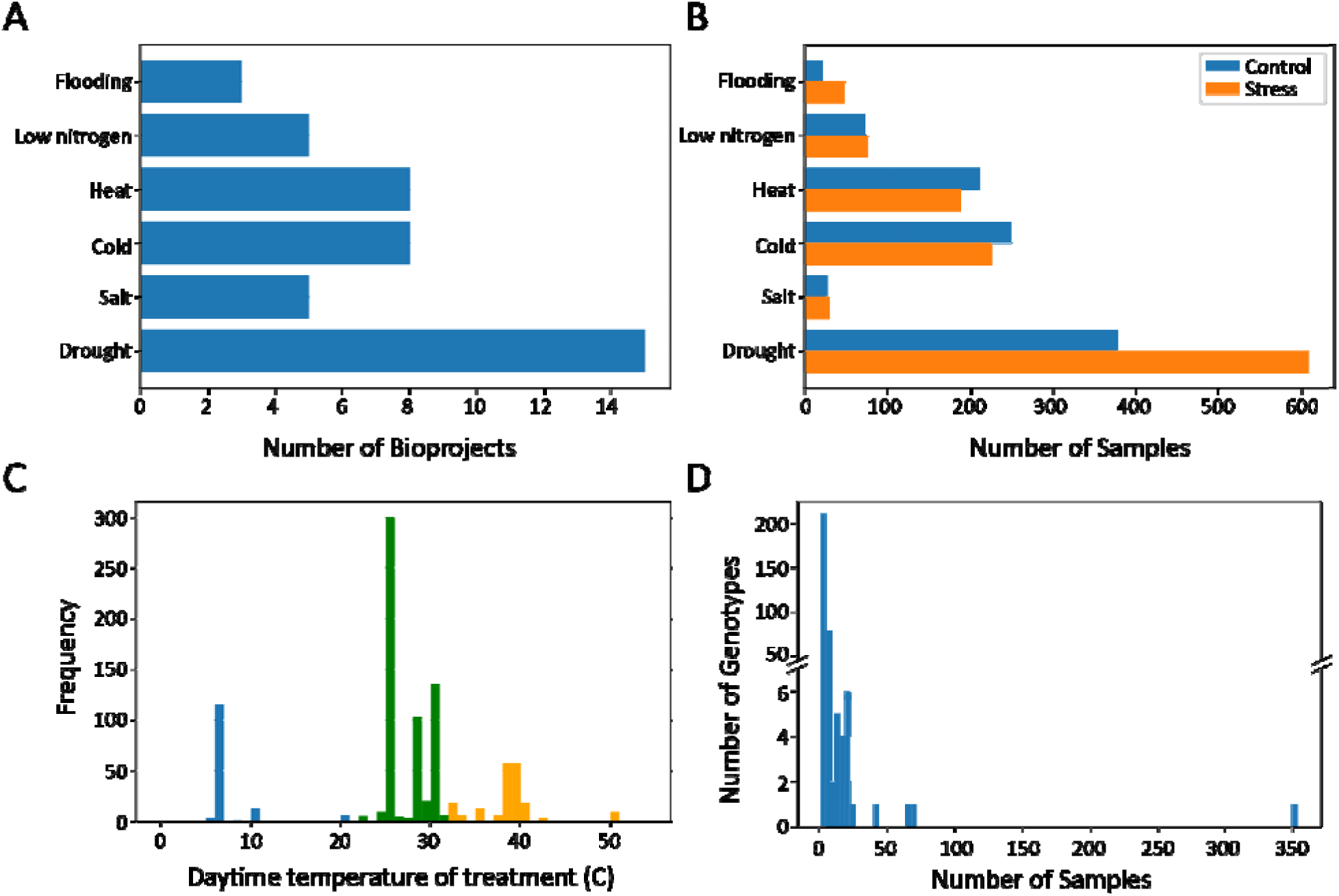
Summary of maize abiotic stress gene expression data. (A) Number of BioProjects per stressor. (B) Number of samples per stressor. (C) Distribution of treatment temperatures, with cold (blue), heat (orange), and control (green) showing distinct means but some overlap at the range edges. (D) Distribution of sample numbers across genotypes, with most genotypes represented by few samples and only a few exceeding 50.

**Table 1:** Numbers of samples for each tissue type in the dataset.

| **Tissue** | **Number of samples** |
| --- | --- |
| Leaf | 1,394 |
| Root | 278 |
| Shoot | 46 |
| Ear | 44 |
| Tassel | 39 |
| Embryo | 24 |
| Pollen | 16 |
| Stalk | 10 |
| Silk | 9 |
| Kernel | 4 |
| Ovary | 4 |
| Leaf meristem | 4 |

All RNA-seq samples were downloaded from the SRA and reanalyzed using a common pipeline with the B73 V5 maize genome as the reference. Although substantial variation in gene content has been observed across maize diversity (Hufford et al., 2021), we used a single reference to enable comparisons across all datasets. We tested the impact of reference genome on read mapping rates for different genotypes. RNAseq reads from the inbreds Oh43 and CML69 had similar mapping rates when aligned to their corresponding de novo reference genomes and the B73 V5 reference genome (Supplemental Table 1). Overall, samples had an average read mapping rate of 76% with most having greater than 60% of reads mapped to B73 transcripts (Supplemental Figure 1).

Principal component analysis was used to visualize sample separation. Approximately 28% of the variance in the full dataset was explained by PC1 and 15% by PC2. We found that samples grouped first by tissue type, with photosynthetic and non-photosynthetic tissues segregating along PC1 and with separation within tissues primarily by BioProject (Figure 2A and B). There was little apparent clustering of samples by treatment (Figure 2D), growth environment (Figure 2C), or developmental stage (Supplemental Figure 2). Any clustering by these factors is likely an artifact of BioProject, as each experiment used different growth conditions, sampled at different developmental stages, and may have applied stress treatments differently.

**Figure 2:**
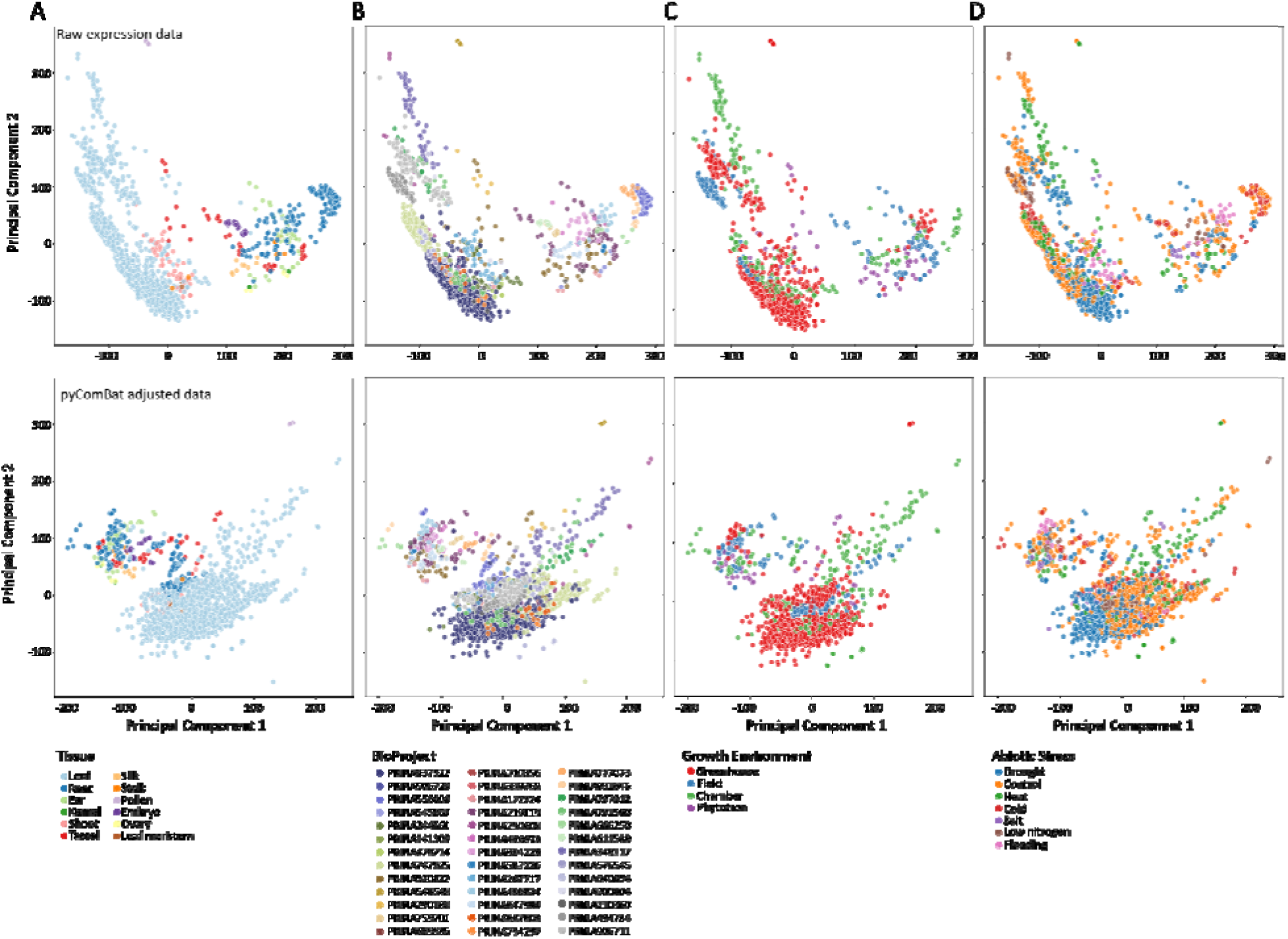
Principal component analysis of maize gene expression data. Principal component analysis (PCA) biplots of log-transformed TPM values, shown without variance correction for BioProject effects (top) and after pyComBat correction (bottom). In the uncorrected data, PC1 explains 28% and PC2 explains 15% of the variance, while in the corrected data, PC1 explains 16% and PC2 explains 14%. Biplots are colored by tissue type (A), BioProject (B), growth environment (C), and treatment (D).

We conducted linear modeling on the first principal component of gene expression, with genotype, BioProject, tissue, and treatment as independent variables. All four independent variables had a highly significant effect on gene expression (p < 0.001). Heterogeneity between genotypes, tissues, treatments, and experiments significantly contributes to the variability in gene expression among samples. Thus, we used pyComBat (Behdenna et al., 2023) to adjust for batch effects and reduce the variance due to BioProject, and re-ran the PCA with the corrected expression values (Figure 2, right column). For the pyComBat adjusted expression, PC1 explained about 16% of variance, and PC2 explained about 14%, and we observed an overall reduction in grouping due to BioProject (Figure 2B). However, some grouping by BioProject was still evident, and this may be due to variability in other factors such as environment or genotype (Figure 2C). We used this pyComBat adjusted expression matrix for all downstream analyses.

Following basic exploration of the dataset, we examined the relative similarity of transcriptomic responses to different abiotic stressors using hierarchical clustering on the batch corrected TPM. For each tissue set, hierarchical clustering identified two main clusters, but these differed substantially in composition between tissue sets (Supplemental Figure 3), likely due to differences in tissue-specific abiotic stress responses.

### Identification and characterization of a core abiotic stress responsive gene set

To identify the core stress-responsive genes across multiple abiotic stressors in maize, we used both set operations of the ratio of gene expression under stress versus normal conditions (fold change or TN-ratios; hereon, set operations) and the top predictive features of random forest based machine learning models. Set operations is the typical method used for identification of core genes in meta-analyses (Cohen and Leach, 2019; Dossa et al., 2019; Tahmasebi et al., 2019; Zhang et al., 2019; Shintani et al., 2024). To capture emergent expression patterns that would be missed from pairwise comparisons, we applied a random forest model to classify stressed samples. In this model, the most important predictive features (i.e., genes) that delineate stressed and healthy samples are defined as core genes. Support vector machine clustering was previously used to identify core stress responsive genes in Arabidopsis (Sanchez-Munoz et al., 2024), but our random forest approach utilizes classification rather than clustering. Given our underlying hypothesis that a core stress transcriptome exists, we expected that a binary random forest classifier model would be able to predict whether a given transcriptome was from a stressed or control sample, even if the model had not been trained on the stressor on which it was being tested. This led to our “hold one stressor out” random forest approach (see methods for more details).

The efficacy of random forest prediction varied across stressors (Supplemental Figure 4), although all area under the ROC curve (AUC) values were greater than 0.5. Salt and drought were consistently predicted most accurately, while low nitrogen, cold, and heat all had AUC values of slightly greater than 0.5. This is similar to the close clustering of temperature stressors with control after hierarchical clustering of treatments (Supplemental Figure 3), which can further be explained by the fact that the low end of control temperatures overlaps with the high end of cold-treatment temperatures, and vice versa for heat (Figure 1C). Thus, some “heat” and “cold”-treated plants may not have been fully physiologically stressed. These results indicate that our machine learning approach was able to capture biological patterns of responses to multiple abiotic stresses. The top features of the model are most predictive of stress vs control samples, and thus, would represent core stress responsive genes.

Using these combined methods, we identified a total of 744 core abiotic stress genes. Table 2 shows a summary of different categories of the core gene sets. Notably, more core genes were found via random forest than by set operations, and more core genes are upregulated than downregulated under stress conditions. The top features in the random forest models were generally differentially expressed under stress compared to control conditions, and there were only 11 core genes that were not differentially expressed.

**Table 2:** Summary of core gene numbers by different methods.

| **Method** | **Number of**<br>**upregulated**<br>**genes** | **Number of**<br>**downregulated**<br>**genes** | **Number of**<br>**genes with no**<br>**differential**<br>**expression** | **Total genes** |
| --- | --- | --- | --- | --- |
| Set operations | 193 | 66 | 0 | 255 |
| Random forest | 462 | 413 | 11 | 524 |
| Combined | 628 | 471 | 11 | 744 |

Once core genes were identified, we tested for overlaps among these genes across these two approaches and whether they were up- or downregulated. We found that genes identified by the same method, whether by random forest or set operations, showed more similarity across different tissue sets than those identified by different methods within the same tissue set (Figure 3A). For this reason, we focused primarily on core and stress-specific genes from all tissues for the rest of this study.

**Figure 3:**
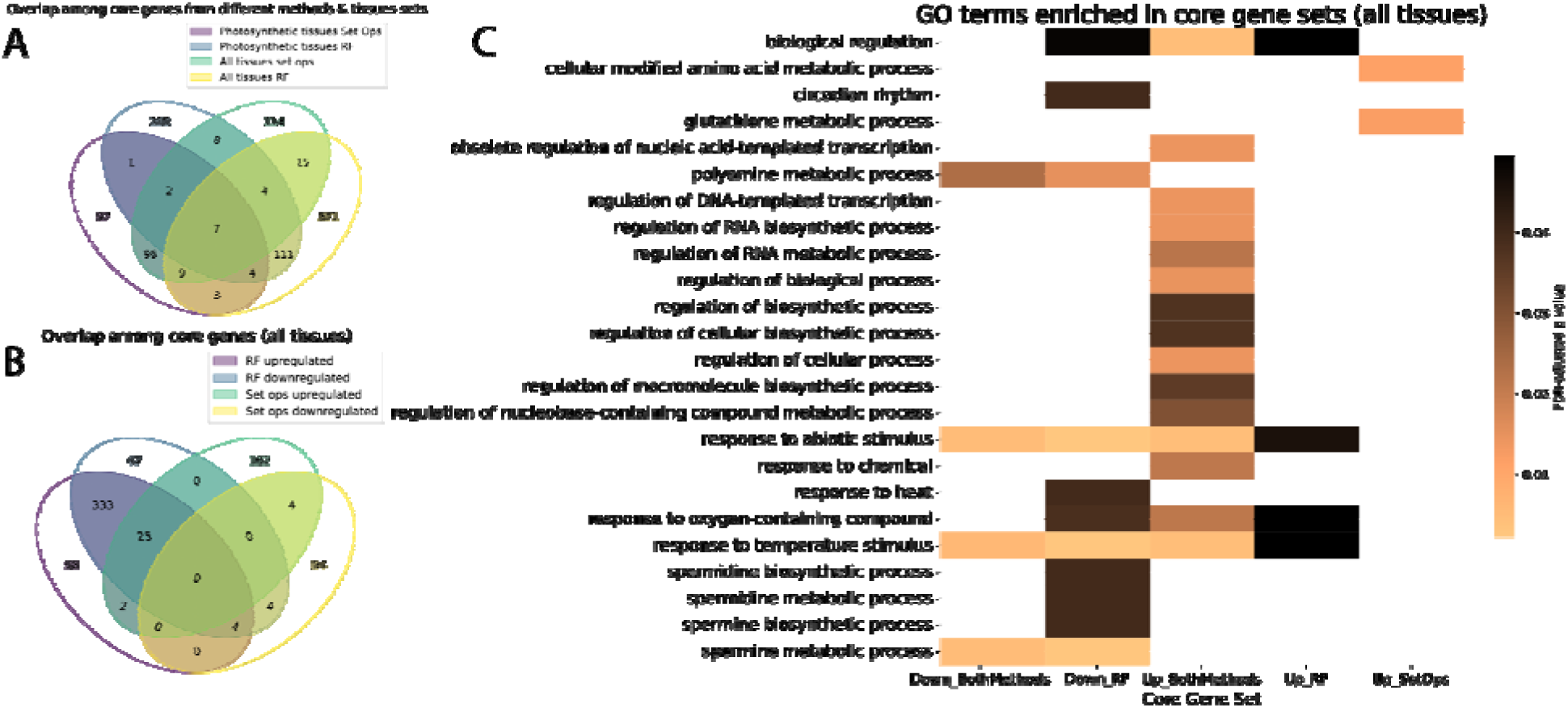
Basic characterization of the core abiotic stress gene sets. (A) Overlap of core gene sets identified by set operations and random forest, as well as overlap between core genes from different tissue sets. (B) Comparison of the overlap between upregulated and downregulated core genes (in any stress condition) for genes identified by random forest and set operations (“set ops”). (C) Gene Ontology (GO) enrichment analysis of core gene sets across tissues, highlighting terms related to regulatory processes, metabolites, and stress response.

Core genes identified by random forest may be differentially expressed in opposite directions for individual stresses, and we observed a large overlap between upregulated and downregulated genes (Figure 3B). This was not the case for core genes from set operations, where by definition the upregulated and downregulated genes were identified separately. There was minimal overlap between core genes identified from different methods (Figure 3B), suggesting that the random forest core genes, identified based on their importance to the models rather than strictly by differential expression, may represent an emergent core response that would not be identified by set operations. Future core stress meta-analyses could benefit from a similar use of machine learning to identify emergent core stress genes rather than simple comparisons.

To investigate the functions of the core abiotic stress genes in maize, we ran GO term enrichment separately on the upregulated and downregulated sets of core genes from each method, including both methods combined. Several enriched terms were found for the core genes (Figure 3C). These were largely distinct between upregulated and downregulated core genes, but four terms, “biological regulation”, “response to oxygen-containing compound”, “response to abiotic stimulus”, and “response to temperature stimulus”, were found across up- and downregulated core gene sets, albeit sometimes with different p-values (Figure 3C). Generic responses to stimuli may take the form of either increases or decreases in abundance, and regulation may be applied by repression or activation.

Notably, GO terms specific to the downregulated core genes include various terms related to polyamine metabolism, especially that of the “higher” polyamines spermidine and spermine (Figure 3C). Polyamines are stress-induced molecules that have protective roles in plants. While the detailed molecular mechanisms underlying this protection remain unclear, plants that overexpress polyamine biosynthetic genes often exhibit enhanced stress tolerance (Minocha et al., 2014; Bano et al., 2020). However, polyamine catabolism can also release reactive oxygen species, so it is possible that polyamines may also contribute to oxidative stress (Minocha et al., 2014). The GO term “circadian rhythm” was also uniquely enriched in downregulated core genes (Figure 3C). Interactions between the circadian clock and abiotic stress responses are complex, but various clock components have been found to be downregulated under different stressors (Sharma et al., 2022). In upregulated core genes, specifically those found via set operations, other metabolism-related GO terms were enriched, including: “cellular modified amino acid metabolic process” and “glutathione metabolic process” (Figure 3C). Amino acids play a crucial role in stress responses, including proline, which functions as a compatible solute (Batista-Silva et al., 2019). Glutathione is also a noted important antioxidant under various stressors (Aslam et al., 2021). In addition, many terms related to transcriptional regulation (i.e. “regulation of RNA biosynthetic process”) were enriched in the upregulated core genes, from both methods combined. This led us to investigate the presence of transcription factors among the core genes.

Stress-specific genes were defined as those that were differentially expressed in response to only one stress condition, in at least one study of that stressor, and were identified by set operations. The number of stress-specific genes varied by stressor (Table 3), with flooding consistently having the fewest stress-specific genes, likely because it also had the fewest experiments (i.e., BioProjects) in the dataset (Figure 1A). Across tissue sets, heat had more upregulated than downregulated stress-specific genes, while drought and cold had more downregulated stress-specific genes (Table 3).

**Table 3:**
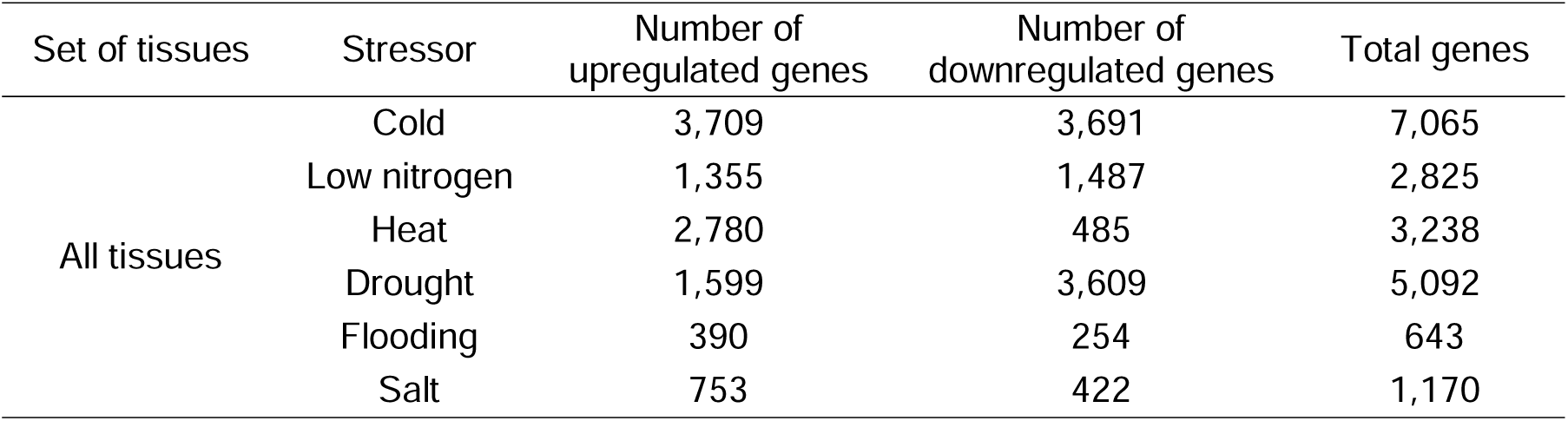
Numbers of stress-specific genes for the six surveyed abiotic stresses in maize.

| Set of tissues | Stressor | Number of upregulated genes | Number of downregulated genes | Total genes |
| --- | --- | --- | --- | --- |
| All tissues | Cold | 3,709 | 3,691 | 7,065 |
|  | Low nitrogen | 1,355 | 1,487 | 2,825 |
|  | Heat | 2,780 | 485 | 3,238 |
|  | Drought | 1,599 | 3,609 | 5,092 |
|  | Flooding | 390 | 254 | 643 |
|  | Salt | 753 | 422 | 1,170 |

There are eleven GO terms that are enriched both in core genes from all tissues and stress-specific genes from all tissues. All of them are related to regulation, including “regulation of DNA-templated transcription”, “regulation of macromolecule biosynthetic process”, and “regulation of biosynthetic process”. These terms were enriched only in downregulated stress-specific genes from cold stress (Supplemental Figure 5), while for core genes, they were enriched in upregulated core genes overall (Figure 3C). Thus, it is possible that the cold stress response in maize is at least partially regulated by upregulated core stress genes.

### Transcription factor enrichment

We used Fisher’s exact test to test for enrichment of transcription factors broadly and specific TF families in sets of core abiotic stress genes. In all cases, upregulated and downregulated core genes were tested separately. For general TF enrichment tests, core gene sets were also separated by method. General TF enrichment was found to be significant for the upregulated and downregulated genes both from RF and from both methods combined (all P<0.001). Significantly enriched and near-enriched TF families were also found (see Table 4). With the exception of the orphan TF family, the enriched and near-enriched TF families have all been previously related to stress tolerance. These include the bZIP family, specifically the ABF subfamily, which is involved in ABA signaling (Yoon et al., 2020); the ERF subfamily of AP2-ERF, which plays a role in flooding response (Mizoi et al., 2012); and the HSFs, which regulate heat shock proteins (HSPs) (Andrási et al., 2021), important molecular chaperones responsive to various abiotic stressors (ul Haq et al., 2019).

**Table 4:** Summary of TF families enriched (adjusted P<0.05) and near-enriched (0.05<P<0.1) in core gene sets.

| Set of tissues | Regulatory direction | TF family | P-value |
| --- | --- | --- | --- |
| All tissues | Upregulated | AP2/ERF-ERF | 0.0468 |
|  |  | C2C2-CO-like | 0.0375 |
|  |  | HSF | 0.005 |
|  |  | Orphans | 0.005 |
|  |  | NAC | 0.0924 |
|  |  | bZIP | 0.0924 |
|  | Downregulated | HSF | 0.0191 |
|  |  | Orphans | 0.007 |

We also ran TF enrichment for the stress-specific genes, treating upregulated and downregulated genes separately. The following stress-specific gene sets were enriched in TFs: upregulated flooding-specific (P=0.0422), upregulated cold-specific (P<0.001), downregulated cold-specific (P<0.001), and downregulated heat-specific (P<0.001). Supplemental Table 2 shows the TF families that were enriched in different stress-specific gene sets. These were largely different from the families enriched in core genes, but there were some families enriched in both core and stress-specific sets. These included AP2/ERF-ERF, which was enriched in upregulated core genes and near-enriched in upregulated heat-specific genes; C2C2-CO-like, which was enriched in upregulated core genes and in upregulated flooding-specific genes; and NAC, which was near-enriched in upregulated core genes and enriched in upregulated heat, downregulated drought, and downregulated flooding-specific genes. ERF and NAC TFs, in particular, are noted for their stress responsiveness, so it is not surprising to find them enriched in both core and stress-specific gene sets.

### Co-expression network analysis of core abiotic stress genes

To further investigate the relationships among core and stress-specific genes and their potential regulatory roles, we constructed a co-expression network, enabling the identification of gene modules, regulatory hubs, and associations between transcription factors and stress-responsive genes. A co-expression network was generated from the batch corrected TPM for all samples using WGCNA (Langfelder and Horvath, 2008; Langfelder and Horvath, 2012). The co-expression network contained a total of 37,216 genes in 23 modules, with a mean module size of 1,618 genes. The largest module (blue) contained 10,959 genes and the smallest (white) contained 103 genes.

Of the 23 co-expression modules, 21 had at least one core gene, and only the darkorange and darkturquoise modules contained no core genes. The five modules with greater than 5% core genes were royalblue (9.6%), green (6.3%), grey60 (6.2%), pink (5.2%), and lightgreen (5.2%). We ran GO term enrichment for these modules to investigate their functions. Enriched GO terms were non-overlapping for four of these five modules (Figure 5B; there were no enriched GO terms found for the green module). The lightgreen module was associated with ethylene response, while the grey60 module was enriched for metabolic processes, particularly nitrogen metabolism. The pink module had the highest number of enriched GO terms, many of which were related to regulation. The royalblue module included terms related to various stimulus and stress responses, as well as protein folding, suggesting the presence of chaperones in this module. A total of 13 co-expression modules contained transcription factors previously identified within the core gene set, and eight of these modules were also enriched in non-TF core genes at P < 0.05P (Fisher’s exact test).

**Figure 4:**
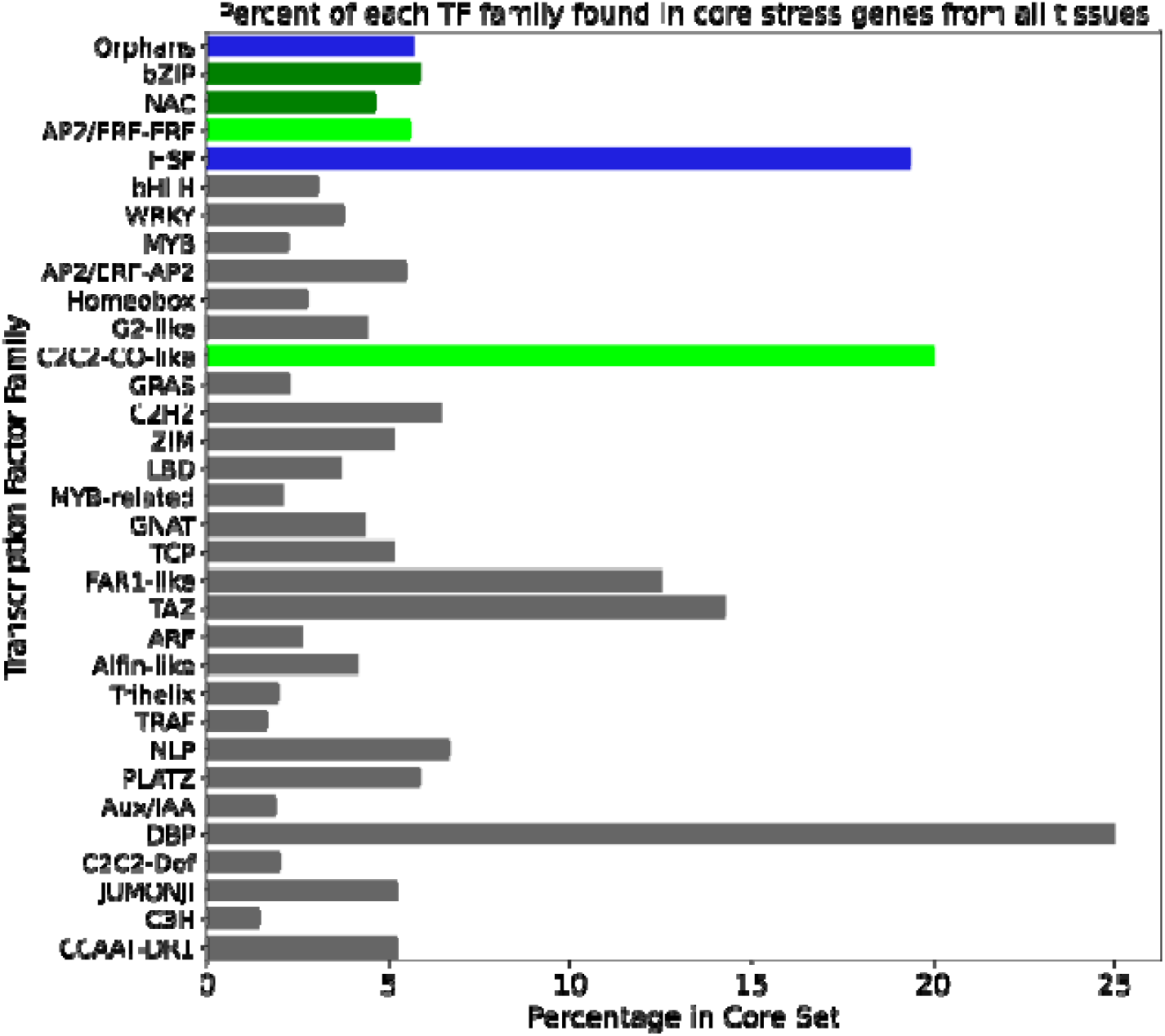
Distribution of transcription factor families in core gene sets across tissues. Light green indicates transcription factor (TF) families significantly enriched in upregulated core genes (p < 0.05, Fisher’s exact test), while dark green denotes families that are near-enriched (0.05 < p < 0.1) in upregulated genes. Blue represents TF families enriched in both upregulated and downregulated core genes. TF families that are not enriched or near-enriched in either core gene set are shown in gray.

**Figure 5:**
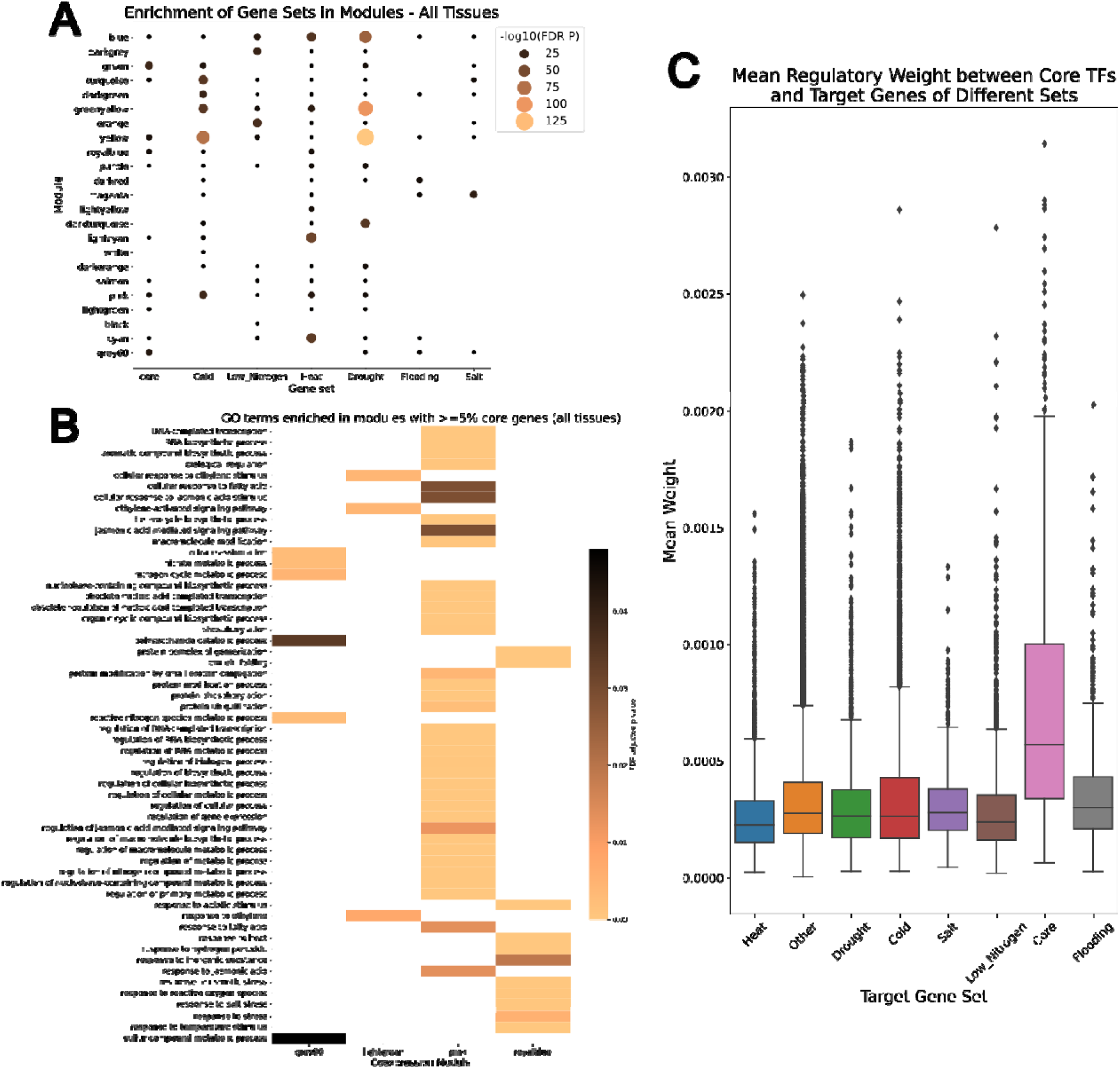
Coexpression Module Analysis of Core and Stress-Specific Gene Sets. (A) Bubble plots showing gene set enrichment in different coexpression modules for core and stress-specific gene sets. The negative log10 of the FDR-adjusted p-value is plotted, with larger bubbles indicating more significant enrichment; non-significant enrichments are not shown. (B) GO term enrichment analysis of modules containing at least 5% core genes. (C) Boxplot of regulatory link weights (mean per target gene) for 33 transcription factors, grouped by different gene sets. In (A), none of the modules enriched in core genes were exclusively enriched in core genes, as all were also enriched in at least one set of stress-specific genes; however, several modules were enriched in all six stress-specific gene sets. In (C), “other” represents genes not present in any of the core or stress-specific gene sets. Core transcription factors from enriched families were significantly more likely to regulate core genes than “other” genes.

Hub genes were identified from the co-expression data using the module membership correlation values, and genes with module membership above the 95th percentile were considered hub genes. Using these criteria, we identified 1,861 hub genes in the maize stress response network. Of these, 14 were core genes, which was not significantly more than expected by chance (Fisher’s exact test, P=0.999). None of the core hub genes were transcription factors. We also tested whether the co-expression modules were enriched in stress-specific genes for each stressor and in core genes using Fisher’s exact test (Figure 5A). In all modules where core genes were enriched, at least some sets of stress-specific genes were also enriched, indicating that core genes are not more strongly coexpressed with each other than with stress-specific genes. The blue, grey60, green, turquoise, lightgreen, royalblue, pink, orange, and greenyellow families all contained at least one core gene TF of one of the enriched or near-enriched families (Table 4), excluding orphans. All these modules except greenyellow and orange were enriched in core genes in all tissues (Figure 5A), and all were enriched in at least 2 stress-specific gene sets. This indicates that the core gene members of the AP2/ERF-ERF, NAC, bZIP, HSF, and C2C2-CO-like TF families are coexpressed with stress-specific stress-specific genes and may regulate these genes.

### Regulatory network analysis of core stress genes

Based on the results of the coexpression network analysis, we hypothesized that core gene transcription factors (TFs) from enriched families regulate not only other core genes but also stress-specific genes. To infer directional regulatory relationships beyond the correlations captured in coexpression networks, we constructed a gene regulatory network (GRN) using GENIE3 (Huynh-Thu et al., 2010). To assess the regulatory influence of core TFs, we used Dunnett’s t-test to compare regulatory link weights between TFs from enriched and near-enriched families and various target gene sets. These included core genes, stress-specific genes for each stress condition, and a background gene set. For each comparison, we generated a distribution of 5,000 p-values from Dunnett’s t-test and calculated the 97.5% confidence interval to determine significance.

We found that the core TFs of interest had significantly higher regulatory weights for other core genes compared to background genes (p=0). In addition, Figure 5C shows that the weights for core gene targets are higher than those for other targets; thus we can conclude that these important core TFs are significantly more likely to regulate core genes than non-core or stress-specific genes. Supplemental Table 3 shows the confidence intervals of p value distributions for the stress-specific versus other targets comparisons. With a 97.5% confidence interval of 0.0039 to 0.0052, there is a significant difference between heat-specific and other target genes; there were no significant differences for any of the other gene set comparisons (Table 5). Figure 5C reveals that on average, the weights for heat-specific target genes were lower than those of other targets. Thus, the important core TFs are significantly less likely to regulate heat-specific genes than the non-core or stress-specific genes.

## DISCUSSION

Abiotic stresses such as drought, flooding, salinity, and temperature extremes trigger similar physiological and biochemical responses in plants. These stresses commonly lead to oxidative stress, cellular damage, and disruptions in metabolic processes, and plants activate a core set of stress-responsive genes to mitigate these disruptions, in addition to the unique and well-characterized pathways for each individual stress. Meta-analyses in various crop and model plant species have found between 20-6,000 core abiotic stress-responsive genes based on overlapping expression profiles and, in one case, support vector machine clustering (Li et al., 2017; Cohen and Leach, 2019; Dossa et al., 2019; Tahmasebi et al., 2019; Zhang et al., 2019; Sanchez-Munoz et al., 2024; Shintani et al., 2024). Previous efforts have focused on a narrow set of abiotic stresses, used microarrays with only partial gene representation, or had sparse gene expression datasets that fail to capture the breadth or diversity of stress responses within a species. Here, we sought to expand on previous work by using a diversity of genotypes, abiotic stresses, tissue types, and stress severity in maize. Using both set operations and random forest classification, we found 744 core genes (Table 2). The number of core genes we found for drought, salt, cold, heat, low nitrogen, and flooding in maize is consistent with previous studies.

We leveraged RNA-seq data from 39 previously published experiments, downloaded from their NCBI BioProjects. Several of the previous meta-analyses analyzed the data from significantly fewer BioProjects than were used in the present study (less than 10 studies each in (Dossa et al., 2019) and (Cohen and Leach, 2019)), and to our knowledge, this is the first such meta-analysis to leverage random forest classification modeling to help identify core genes.

Most previous studies have used only set operations on differentially expressed genes, which is a limited approach in that it defines core genes by their expression under stress alone. In contrast, our random forest approach identified many genes that, while differentially expressed, were not identified as core genes by the set operations method, and in fact identified a few genes that were not differentially expressed under any condition. There was little overlap between the results from the two methods. This indicates that the random forest method picked up on emergent aspects of core stress response, broadening the core stress gene set. This new methodology may be informative for future core stress meta-analyses.

Gene expression changes under abiotic stress are achieved through the action of regulators such as transcription factors (TFs). We found that the core stress gene set was enriched in TFs, including particular families such as heat shock factors (HSFs), ethylene response factors (AP2/ERF-ERFs), C2C2-CO-like, NAC, and bZIP (see Table 4). These families have well-characterized roles in abiotic stresses in plants (Mizoi et al., 2012; Yoon et al., 2020; Andrási et al., 2021). For instance, the heat shock factors (HSF) are the direct regulators of heat shock proteins (HSPs) (Andrási et al., 2021), which act as molecular chaperones under various abiotic stress conditions (ul Haq et al., 2019). Core gene members of the HSF family are present in a co-expression module with enriched GO terms related to “protein folding”, and are thus likely involved in regulating HSPs and thus protein stability under all abiotic stresses.

The members of the AP2/ERF-ERF TF family have been found to be responsive to multiple abiotic stresses, including flooding, drought, and salt (Mizoi et al., 2012; Debbarma et al., 2019). At least one member of this family, *TaERF4* from wheat, is likely a negative regulator of salt response (Dong et al., 2012). Furthermore, gene editing of members of this family has yielded improved stress tolerance in multiple cases (Debbarma et al., 2019). The AP2/ERF-ERFs were enriched in upregulated core genes as well as upregulated heat stress-specific genes for all tissues (Table 4), suggesting that they are important regulators of both core stress and heat-specific responses in maize. Core gene members of this family are found in a co-expression module where the only enriched GO terms are all related to ethylene (Figure 5B), highlighting a potential regulation by ethylene.

All TF families enriched or near-enriched in core genes were present in modules that were also enriched in at least two stress-specific gene sets (Figure 5A). This suggests that, given their coexpression with both other core genes (in some modules) and stress-specific genes, these core TFs may regulate both the core abiotic stress response and stress-specific responses. However, further analysis using a gene regulatory network reveals that while these TFs are significantly more likely to regulate core genes than genes that are neither core nor stress-specific, they do not show a similar pattern for any stress-specific gene set (Figure 5C). Notably, these TFs are significantly less likely to regulate heat-specific genes compared to other sets (Figure 5C). This indicates that, although some crosstalk may exist between the regulation of core and stress-specific gene sets, distinct regulatory pathways likely govern the different sets of genes.

Thus, these core stress TFs may make good targets for improvement of abiotic stress tolerance at least in maize, and possibly in other species as well, especially considering that overexpression of stress-involved TFs has previously led to improved stress tolerance in many cases (Nakashima et al., 2012; Hoang et al., 2014; Baillo et al., 2019). However, further studies are required to elucidate the molecular mechanisms of the maize core stress genes identified in this study. For example, if a TF acts as a repressor on genes that would be beneficial if expressed under stress, overexpression of that TF could lead to a reduction in stress tolerance rather than the desired increase. This omics-only meta-analysis does not provide enough information to say definitively whether the core genes would be good targets for crop improvement, but we hope that follow-up studies will provide more information.

## ACKNOWLEDGEMENTS

This work is supported by NSF Grant MCBL1817347 to R.V., by the NSF Research Traineeship Program (DGE-1828149) to A.C.H.P., and by predoctoral training award T32-GM110523 from the National Institute of General Medical Sciences of the NIH to J.D.P. The authors would like to thank Dr. Scott Pardo for statistical consultation, Dr. Addie Thompson and Brandon Webster for providing data ahead of publication, and Dr. Serena Lotreck for assistance with random forest methods.

## AUTHOR CONTRIBUTIONS

A.C.H.P., J.D.P., and R.V. designed the research; A.C.H.P. performed the research and analyzed the data; A.C.H.P. wrote the manuscript; J.D.P. and R.V. reviewed and edited the manuscript.

