## Supplemental Figures/Tables for "Stress-responsive transcription factor families are key components of the core abiotic stress response in maize"

SUPPLEMENTAL FIGURES AND TABLES

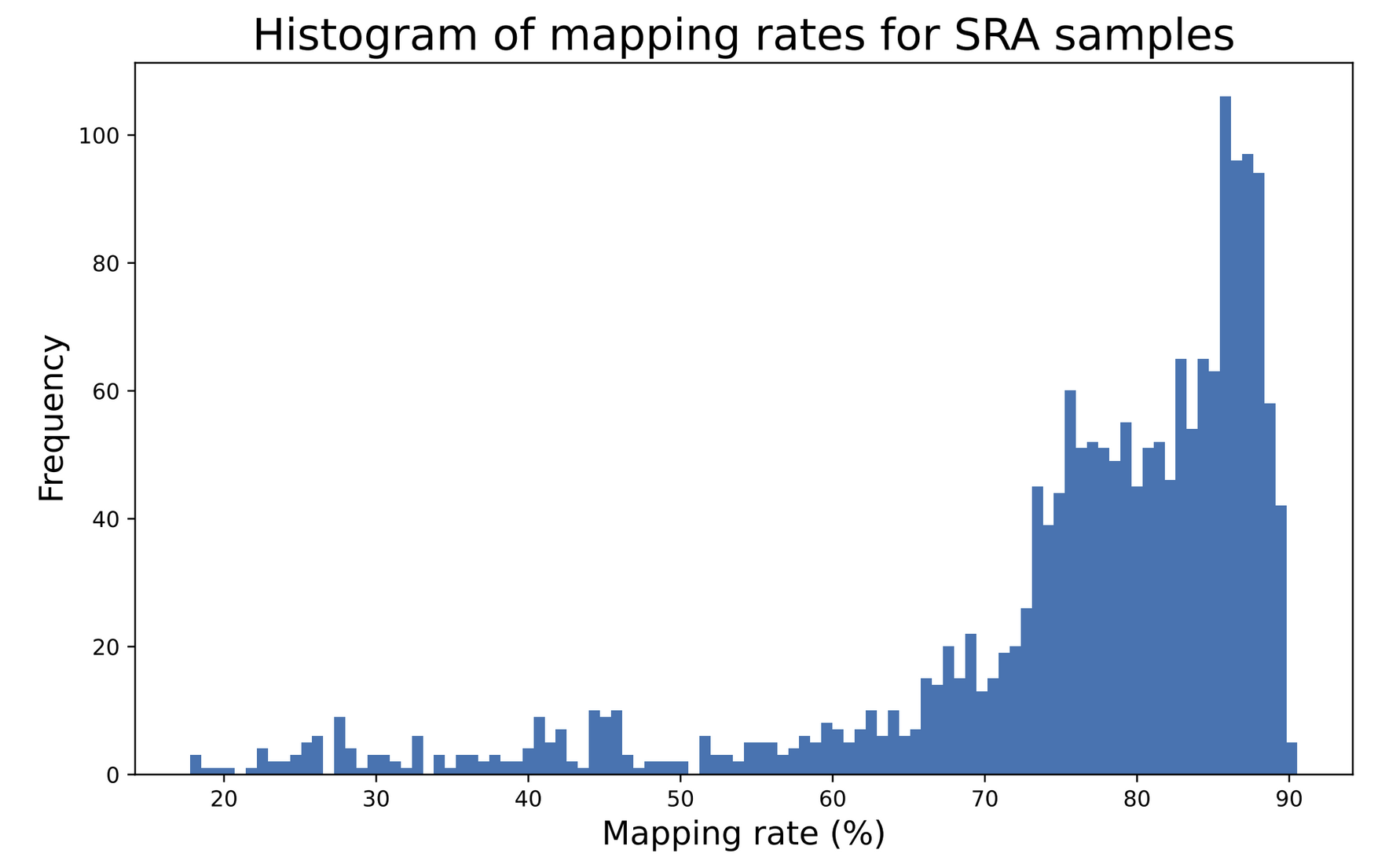

**Supplemental Figure 1: Salmon pseudoalignment mapping rates for all samples downloaded from the NCBI Sequence Read Archive and processed against the B73 V5 genome.**

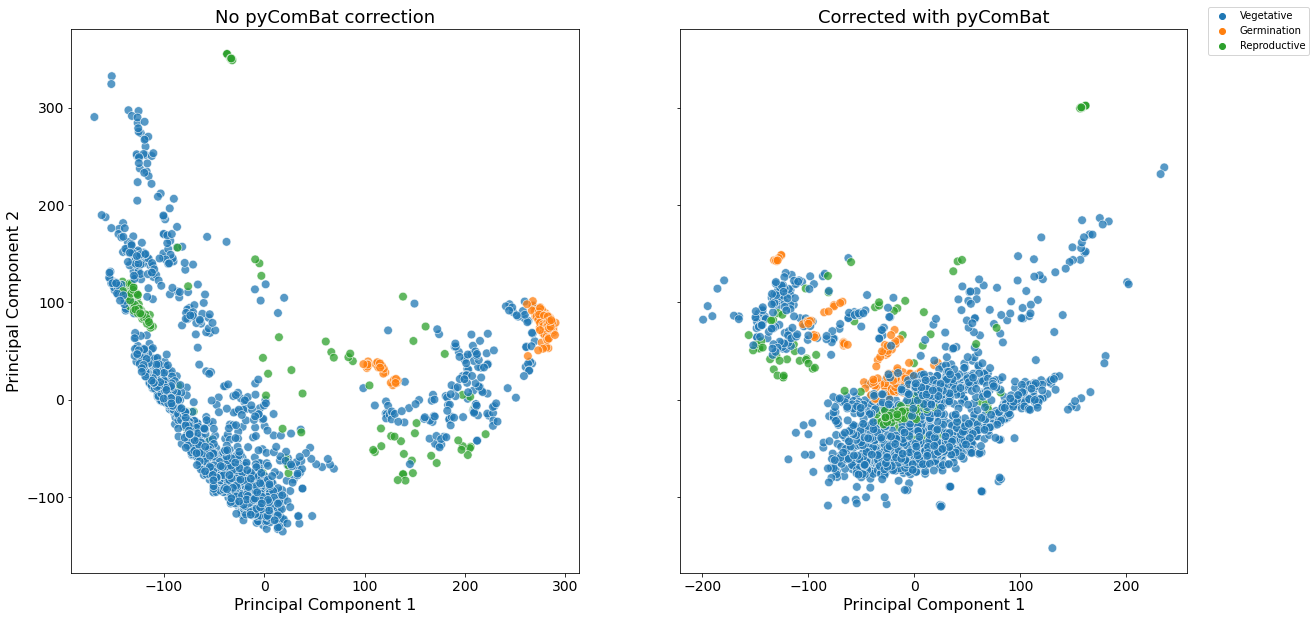

**Supplemental Figure 2: Principal component analysis biplot colored by developmental stage (vegetative, reproductive, or germination).** Uncorrected log(TPM) are on the left, corrected on the right.

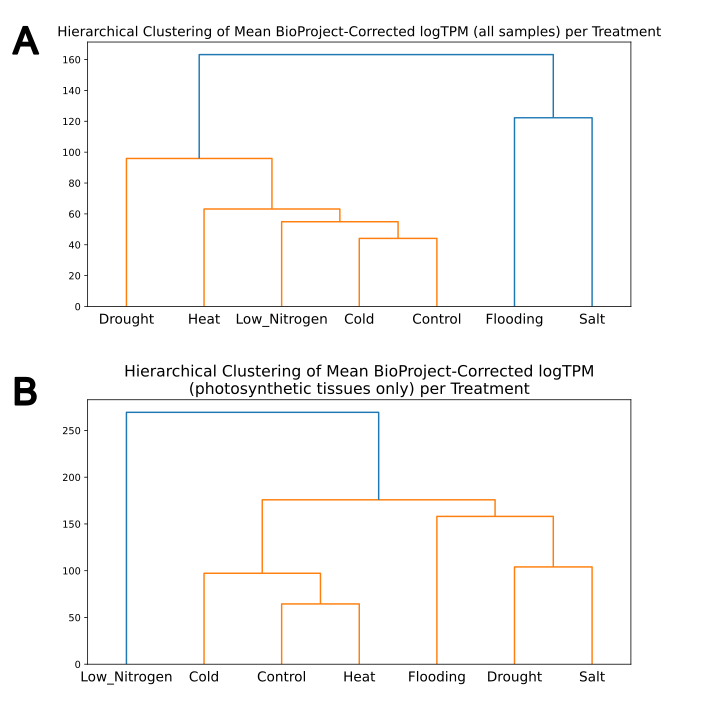

**Supplemental Figure 3: Hierarchical clustering of transcriptomes for different treatments, for all tissues (A) and photosynthetic tissues only (B).**

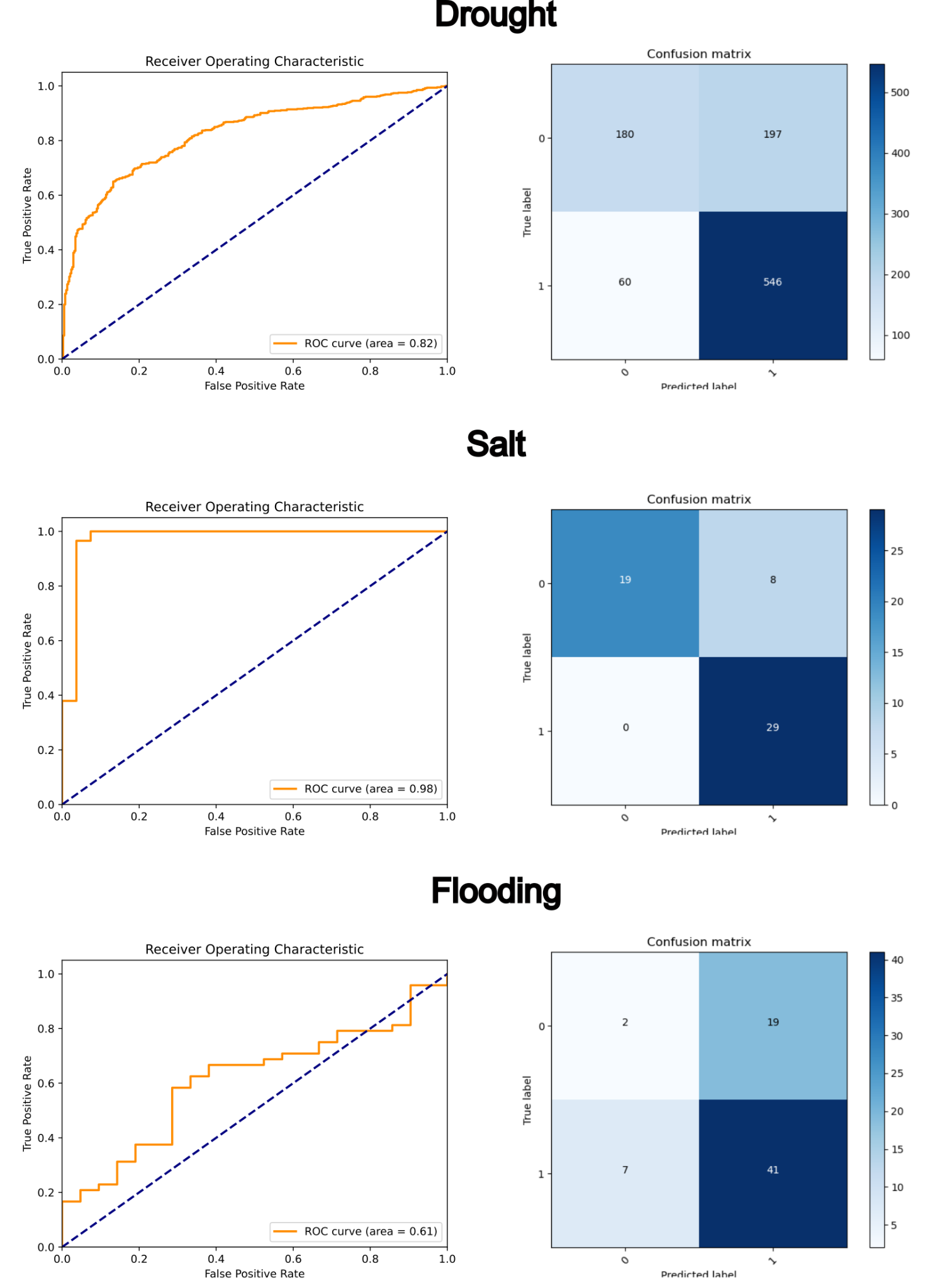

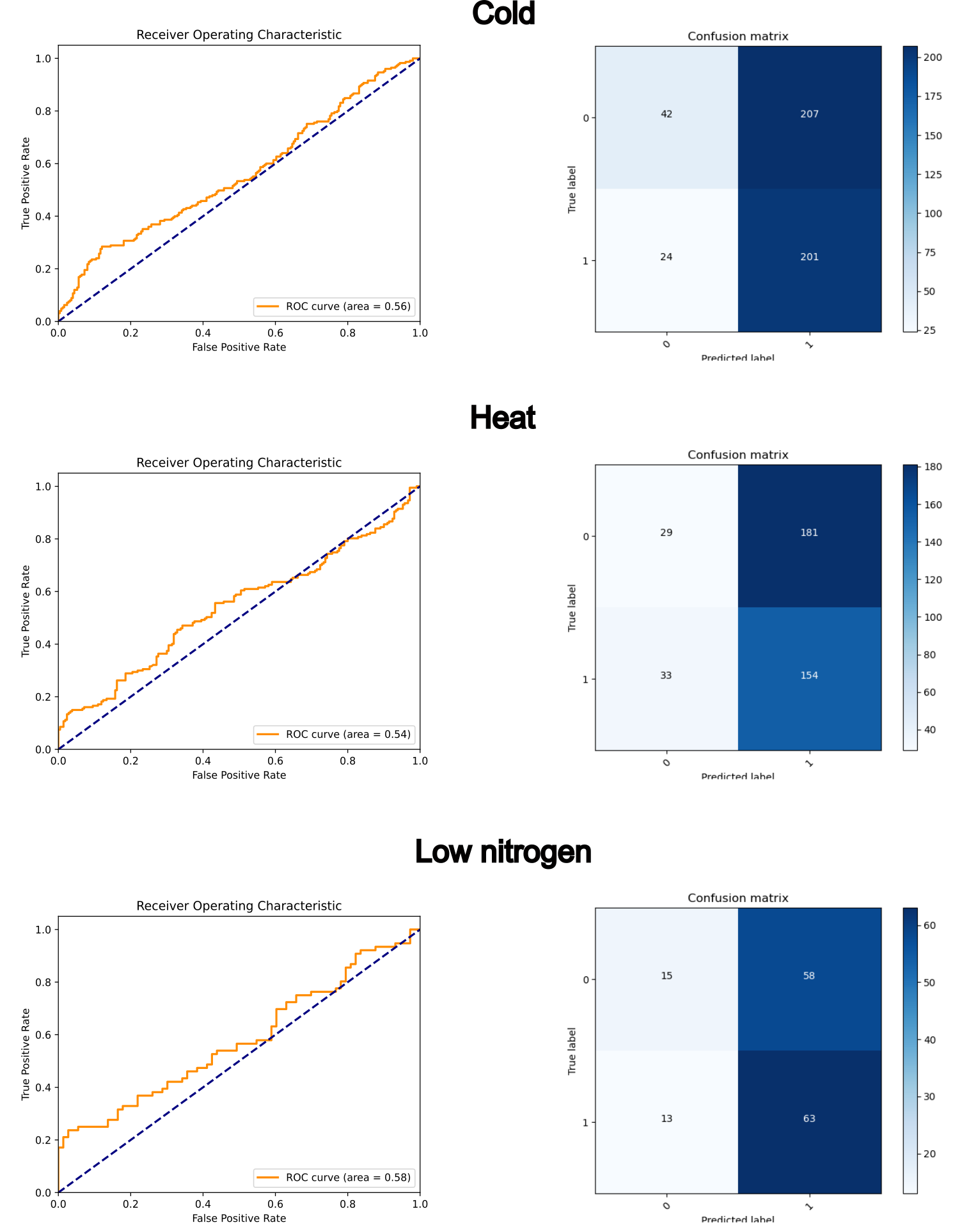

**Supplemental Figure 4: Model performance assessment for random forest classification.** Receiver operating characteristic (ROC) curves and confusion matrices for random forest classification models, with each stressor’s data held out as the test set.

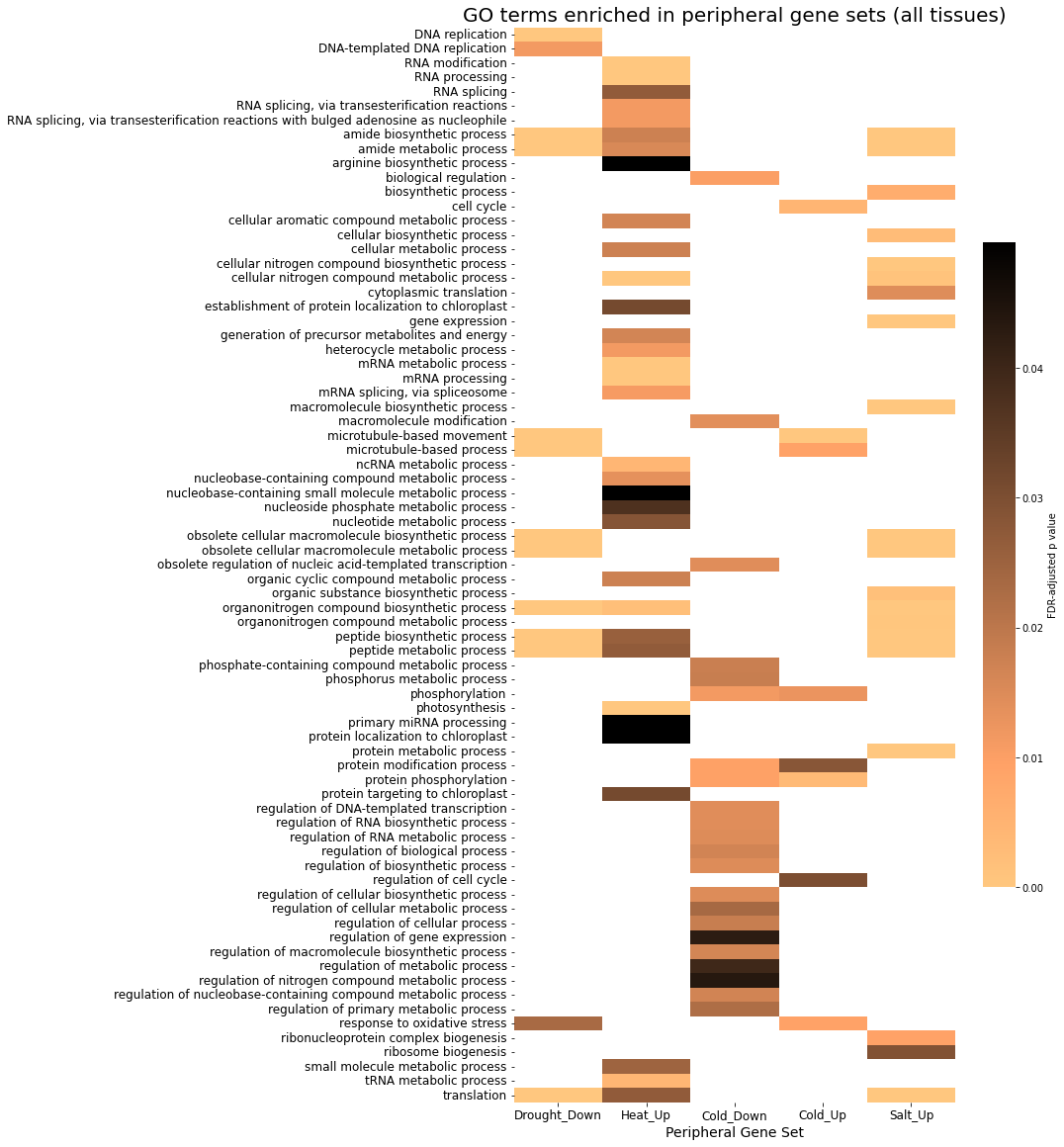

**Supplemental Figure 5: Enriched GO terms in stress-specific gene sets for all tissues.** Lighter colors indicate higher enrichment. Up, upregulated; Down, downregulated. Only the five stress-specific gene sets shown had any GO term enrichment.

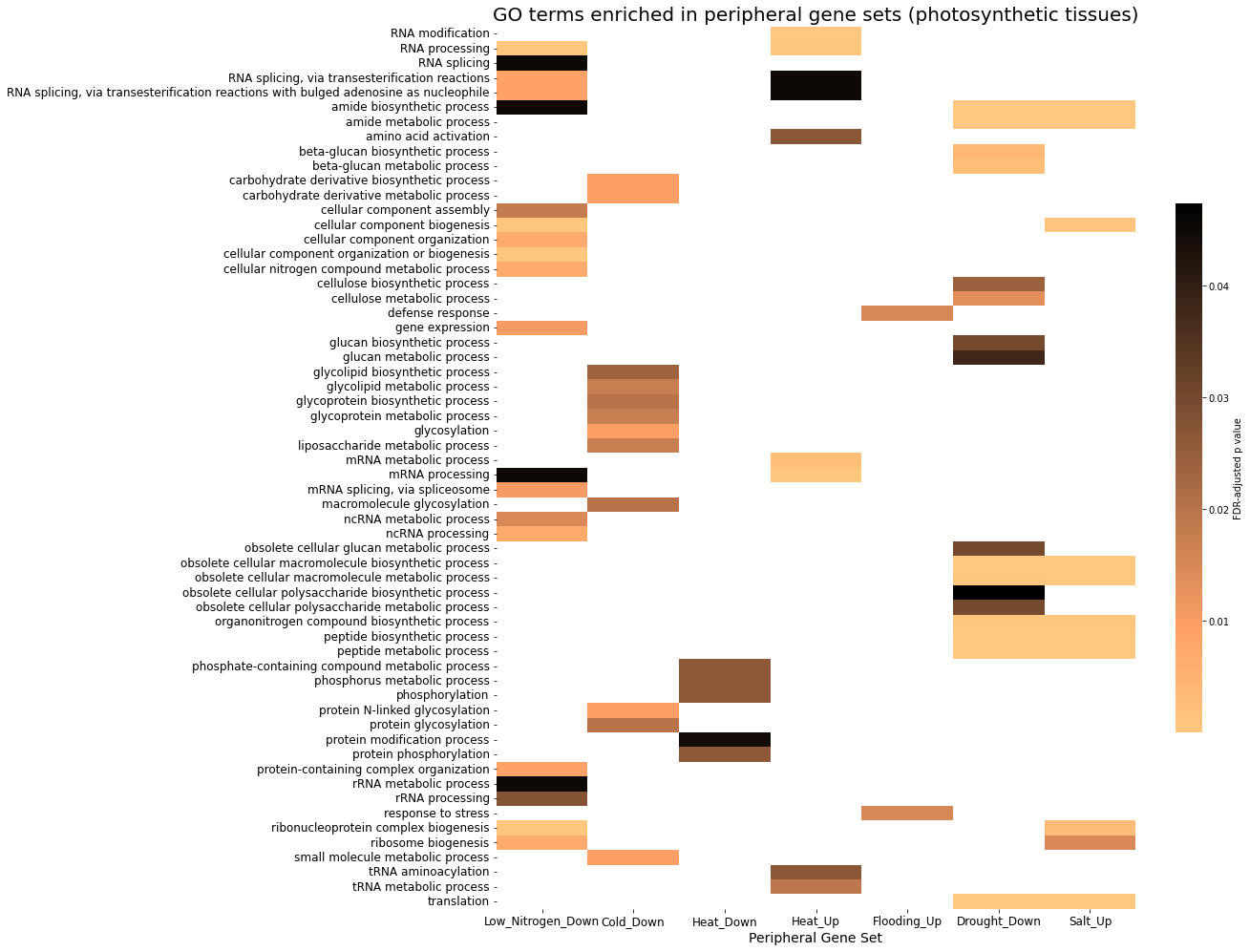

**Supplemental Figure 6: Enriched GO terms in stress-specific genes from photosynthetic tissues only.** A lighter color indicates higher enrichment (lower P-value), except white which indicates no enrichment. Only the gene sets listed on the X axis had any enriched GO terms.

**Supplemental Table 1: Salmon mapping rates of two genotypes of maize when mapped to their own de novo reference genomes and to the B73 v5 reference.**

| Genotype | Mean mapping rate against self | Mean mapping rate against B73 |
| --- | --- | --- |
| Oh43 | 89.08% | 85.45% |
| CML69 | 83.86% | 79.62% |

**Supplemental Table 2: TF families enriched in different stress-specific stress gene sets. P-values given are from Fisher’s exact test, corrected with FDR.**

| Set of tissues | Stressor | Regulatory direction | TF family | P-value |
| --- | --- | --- | --- | --- |
| All tissues | Cold | Upregulated | HMG | 0.0424 |
|  |  |  | OVATE | 0.0424 |
|  |  |  | SHI/STY (SRS) | 0.0094 |
|  |  | Downregulated | C2H2 | 0.0749 |
|  |  |  | WRKY | 0.0749 |
|  | Low nitrogen | Upregulated | CCAAT-HAP2 | 0.0619 |
|  |  |  | ZF-HD | 0.0234 |
|  | Heat | Upregulated | SWI/SNF-SWI3 | 0.0022 |
|  |  |  | mTERF | 0.0775 |
|  |  | Downregulated | ARF | <0.0001 |
|  |  |  | CCAAT-HAP2 | 0.0012 |
|  |  |  | SBP | 0.002 |
|  |  |  | Sigma70-like | 0.0661 |
|  | Drought | Upregulated | C2C2-YABBY | 0.0909 |
|  |  | Downregulated | CCAAT-DR1 | 0.0623 |
|  | Flooding | Upregulated | C2C2-CO-like | 0.0283 |
|  |  | Downregulated | MYB | 0.0087 |
|  |  |  | NAC | 0.0385 |

**Supplemental Table 3: Results of statistical analysis with Dunnett’s t test on gene regulatory network weights.** Each gene set listed here was compared to other target genes not belonging to any of those gene sets. Regulatory links were only considered between 33 transcription factors in the core gene set, belonging to enriched or near-enriched families (Figure 5), and target genes. Confidence intervals of 97.5% were calculated for distributions of 5,000 Dunnett’s t p values. Core genes are not shown; confidence intervals could not be calculated because all p values were 0.

| Gene Set | Lower confidence interval | Upper confidence interval |
| --- | --- | --- |
| Heat | 0.003935385 | 0.005192344 |
| Low nitrogen | 0.09833538 | 0.11006542 |
| Drought | 0.221322 | 0.2385571 |
| Salt | 0.3975272 | 0.4182547 |
| Flooding | 0.665957 | 0.6872534 |
| Cold | 0.8249245 | 0.8410558 |
